# PolyGenie: a reproducible Nextflow pipeline for phenome-wide association studies using polygenic risk scores

**DOI:** 10.1101/2025.09.05.674501

**Authors:** Xavier Farré, Mireia Gasco, Natalia Blay, Rafael de Cid

## Abstract

**Summary:** Phenome-wide association studies (PheWAS) using polygenic risk scores (PRS) offer a powerful framework for exploring the shared genetic architecture of complex traits across diverse phenotypic domains. However, no standardized, portable pipeline exists to facilitate their systematic execution and visualization in arbitrary population cohorts. We present PolyGenie, an open-source Nextflow pipeline that takes precomputed PRS and cohort phenotype data as input and performs scalable PheWAS analysis across binary and continuous outcomes. The pipeline produces regression results and percentile-based prevalence estimates, which are stored in a SQLite database and visualized through an interactive Dash web application. To demonstrate its utility and reproducibility, we provide a fully worked example using the GCAT cohort, applying 135 PRS to a broad range of clinical, molecular, and lifestyle phenotypes. PolyGenie is designed to be deployed on any cohort with minimal configuration, enabling standardized cross-trait analyses and interactive exploration of genetic risk.

**Availability and Implementation:** PolyGenie is freely available at https://github.com/gcatbiobank/polygenie-pipeline. The GCAT implementation can be accessed at https://polygenie.igtp.cat, and the corresponding adapted Dash source code is available at https://github.com/gcatbiobank/polygenie-gcat.

**Supplementary information:** Supplementary data are available at Bioinformatics online.

## Introduction

Genome-wide association studies (GWAS) have transformed our understanding of the genetic basis of complex traits and diseases by identifying thousands of associated loci (1). However, the highly polygenic nature of these traits, characterised by contributions from numerous genetic variants with small effect sizes, makes it difficult to use individual variants for accurate risk prediction in common disorders.

Polygenic risk scores (PRS) have emerged as powerful tools for quantifying genetic susceptibility to complex traits and diseases by aggregating the effects of numerous genetic variants (2). PRS not only enhance the prediction of an individual’s risk for specific traits but also enables researchers to explore genetic associations beyond a single trait. By examining how the polygenic risk score (PRS) for a given trait associates with a broad range of other phenotypes and diseases, a phenome-wide association study (PheWAS), researchers can uncover shared genetic architecture and pleiotropic effects, offering insight into the biological mechanisms that connect seemingly distinct conditions.

The growing availability of GWAS summary statistics, along with modern PRS techniques that go beyond selecting GWAS significant hits, and pipelines that ease their use (3), offer a powerful opportunity to deepen our understanding of the interplay between genetic susceptibility to complex traits and diseases and the wide array of variables captured in population cohorts. Particularly, population cohorts linked to clinical records and enriched with multi-layer molecular data (e.g., genomics, metabolomics, exposomics) provide exceptional power to uncover biologically meaningful associations and translate genetic insights into actionable knowledge. However, systematic execution of these analyses requires handling heterogeneous phenotype data across multiple categories, applying appropriate regression models, and presenting results in a way to support rapid hypothesis generation. These steps are rarely standardized, making analyses difficult to reproduce or adapt to new cohorts.

To this end we present PolyGenie, an interactive web-based resource that enables exploration of associations between PRS for 135 traits and a wide array of phenotypes, including diseases, metabolite levels, and lifestyle variables, based on data from the GCAT cohort (4), a large population-based cohort from Southern Europe. PolyGenie is set to expand its coverage to the entire GCAT cohort as additional genotype data and new incident cases become available, with dynamic medical updates planned annually, enhancing the platform’s analytical capacity and overall impact. Additionally, PolyGenie supports exploration by providing intuitive visualisations that facilitate hypothesis generation. By deepening our understanding of the genetic basis of health and disease, the platform empowers future research and contributes to informed public health decision-making.

Several tools support large-scale PRS and PheWAS analyses, each serving distinct purposes. The PRS Atlas (5) and SpPGS (6) are valuable population-level references for linking polygenic scores to traits in UK and Spanish populations, but are inherently static resources tied to specific cohorts and methods, making them unsuitable for deployment on new cohorts or phenotype sets. The R PheWAS package (7) and Python-based tools such as pyPheWAS (8) offer analytical flexibility, but require command-line expertise and lack integrated pipelines or interactive visualization, imposing significant burden when scaling to hundreds of PRS-phenotype combinations. Specialized visualization platforms such as PheWAS-ME (9) and pyPheWAS Explorer (10) provide rich interactive dashboards for exploring individual-level associations and multimorbidity patterns, but operate as post-hoc visualization layers that require prior completion of separate statistical analyses elsewhere. The PheWAS Catalog (11) serves as a valuable reference for previously published PheWAS results but does not support redeployment to new cohorts or custom PRS inputs.

Here we present PolyGenie, an open-source pipeline that addresses this gap by focusing on the PheWAS analysis and visualization steps, accepting precomputed PRS as input. This design decision deliberately separates PRS generation, which users can perform with any method of their choice (e.g., MegaPRS (12), PRSice-2 (13), PRScs (14)), from the standardized downstream analysis, making the pipeline applicable to any cohort. PolyGenie is implemented in Nextflow (15), ensuring portability across computing environments, and ships with a fully configured example based on the GCAT cohort that serves both as a working demonstration and as a reproducibility resource.

## Methods

### Pipeline design and inputs

PolyGenie employs a hierarchical configuration structure that separates user configuration from data specification. A single YAML configuration file serves as the entry point, specifying paths to metadata files and controlling analysis parameters. Two metadata CSV files (one for PRS, one for phenotypes) then direct the pipeline to the actual data files and define their properties: file paths, variable types, analysis-specific settings such as sex stratification, and phenotype-specific covariates. The actual data inputs consist of (i) precomputed PRS files, one per trait, in standard tab-delimited format; (ii) phenotype data files organized by category (e.g., ICD-10 codes, metabolites, questionnaire variables); and (iii) a covariate file containing individual-level variables for regression adjustment.

This metadata-driven design means that adapting the pipeline to a new cohort requires only preparing the input files in the specified formats and editing the configuration, with no changes to pipeline code. Detailed format specifications and a worked example are provided in the repository documentation.

### Pipeline workflow

The pipeline is implemented in Nextflow DSL2 and comprises two stages. In the preprocessing stage, CHECK_PRS_FILES validates each PRS file against its metadata entry, verifying file existence and expected column structure. CHECK_PHENOTYPE_FILES validates each phenotype variable, filtering out binary traits that fall below a user-defined minimum case threshold (default: 10 cases). Both steps produce filtered metadata files and log reports that are archived in the output directory.

In the analysis stage, COMPUTE_PRS_PERCENTILES calculates the prevalence or mean value of each phenotype across PRS percentile bins (configurable, default: 100 bins), producing per-PRS output files suitable for prevalence-by-percentile visualization. COMPUTE_PRS_REGRESSIONS runs linear or logistic regression for each PRS-phenotype pair, comparing the top quantile group against the bottom group or the rest of the cohort (configurable). Multiple regression configurations, varying the number of quantile groups and the reference category definition, can be run simultaneously by specifying multiple entries in the configuration file. Both analysis processes run in parallel across PRS, scaling naturally on HPC clusters and cloud environments via Nextflow’s executor abstraction.

Regression models are implemented in Python using statsmodels, with parallelization across phenotypes via joblib. PRS values are optionally z-score normalized before analysis. Covariates are specified globally in the configuration and can be supplemented with phenotype-specific covariates in the phenotype metadata file.

### Database and web application

After the pipeline completes, results are ingested into a SQLite database using a provided loader script, which populates tables for regression results, percentile data, PRS metadata, and GWAS source information. The web application is implemented in Plotly Dash and queries this database to render three primary views: (i) a PheWAS scatter plot displaying signed log_10_(p-value) across phenotypic domains for a selected PRS; (ii) a prevalence-by-percentile plot for a selected PRS–phenotype pair, stratified by sex; and (iii) a top-hits table with exportable results. The application is containerized and can be deployed locally or on a server with minimal configuration.

### GCAT cohort implementation

To demonstrate the pipeline and provide a reproducibility resource, we applied PolyGenie to the GCAT cohort, a comprehensive population-based living cohort of approximately 20,000 middle-aged individuals (40–65 years old) from Catalonia (4). Recruitment was conducted between 2014 and 2018, and participants are under ongoing active follow-up through continuous linkage to Electronic Health Records (EHRs) from Catalonia’s public healthcare system, providing access to over 15 years of longitudinal clinical data that are regularly updated over time. In addition, genotype data are available, enabling integrated dynamic analyses of genetic, clinical and environmental determinants of health and disease (http://igtp.cat/gcat).

In this study, we utilised available genetic data from a subset (n∼5,000) of the GCAT cohort. Genome-wide genotyping was performed using the Infinium Expanded Multi-Ethnic Genotyping Array (MEGAEx) (Illumina, San Diego, California, USA). All included participants were of Iberian descent with White-Western European ancestry, as determined by self-reported data and principal component analysis (PCA) (16). Imputation was carried out using the TOPMed reference panel (17), resulting in approximately 20 million unique autosomal variants. Only GCAT participants who passed stringent quality control measures were included (N=4,988). Variants with a minor allele frequency (MAF) greater than 0.001 and an imputation quality score (R^2^) above 0.3 were retained for downstream analyses (N=17,203,152). Genotype data are accessible through the European Genome-phenome Archive (EGA) under accession ID EGAD00010001664.

PRS for 135 traits were computed externally using the GenoPred pipeline (18) and MegaPRS method (12) applied to publicly available GWAS summary statistics (Supplementary Table 1), and provided as input to PolyGenie. Phenotypes included diseases coded using ICD-10 and Phecodes (19), metabolomics measurements, and questionnaire variables were quality-filtered to 1,483 traits, resulting in over 200,000 PRS-phenotype associations tested. All regression models were adjusted for age, sex, and the first ten principal components. Metabolite models additionally included a covariate for chylomicrons to account for non-fasting conditions. A Bonferroni correction was applied within each phenotype category independently.

### User interface and visualisation

PolyGenie offers a user-friendly, interactive interface designed to facilitate the exploration of phenome-wide associations for selected polygenic risk scores (PRS). Users begin by selecting a PRS of interest from a searchable dropdown list. They can then choose to compare individuals in the top decile or quartile of the PRS distribution with those in the bottom decile or the rest of the cohort.

The primary visualisation is a PheWAS-style scatter plot, where outcomes are grouped by domain along the x-axis. The y-axis represents the signed log_10_(p-value), allowing users to assess both the strength and direction of associations. Hover functionality provides detailed summary statistics for each point, including effect sizes and p-values. Information about the GWAS source used to construct each PRS is also displayed below the main plot.

Upon selecting a specific phenotype, users are presented with a secondary plot showing the prevalence of that condition across PRS percentiles. This visualisation is accompanied by a data table summarizing the number of affected individuals in the cohort.

Beyond graphical exploration, the GCAT implementation demonstrates how the Dash interface can be customized to support multiple analytical workflows. In this instance, results can be exported in tabular format for external analysis, integrated reference tabs provide searchable GWAS source information and trait metadata, and cohort-level summaries (age distribution, BMI) contextualize genetic findings within the study population. However, these interface features are customizable, users deploying PolyGenie to new cohorts can adapt the web application layout, metrics, and metadata displays to match their specific study design and analytical needs, leveraging the provided Dash source code as a starting template.

### Applying PolyGenie to Explore Sex Differences in Polygenic Risk for Frailty-Associated Traits

To illustrate the utility of PolyGenie’s prevalence-by-percentile visualization, we examined associations between the Fried frailty polygenic risk score, derived from a genome-wide association study of frailty (20), and two clinically relevant outcomes in the GCAT cohort: overweight/obesity (ICD-10: E66) and major depressive disorder (ICD-10: F32). Both outcomes have established epidemiological and genetic links to frailty through shared behavioral, physiological, and social pathways (21).

PolyGenie’s interactive prevalence plots revealed a clear dose–response relationship between higher frailty PRS and increased prevalence of both outcomes (Figures 2a and 2b). For overweight and obesity, prevalence increased approximately linearly across PRS percentiles, with comparable trends in males and females. For major depressive disorder, a similar gradient was observed, but with substantially higher prevalence in females across all percentiles — consistent with the approximately twofold higher 12-month MDD prevalence in women compared to men reported in epidemiological literature (22), and demonstrating the tool’s sensitivity to both biological and socially mediated sex differences. These patterns align with prior GWAS findings reporting genetic correlations between frailty, adiposity, and neuropsychiatric traits (20), and are consistent with a bidirectional interplay among these aging-related syndromes (23).

**Figure 1.**
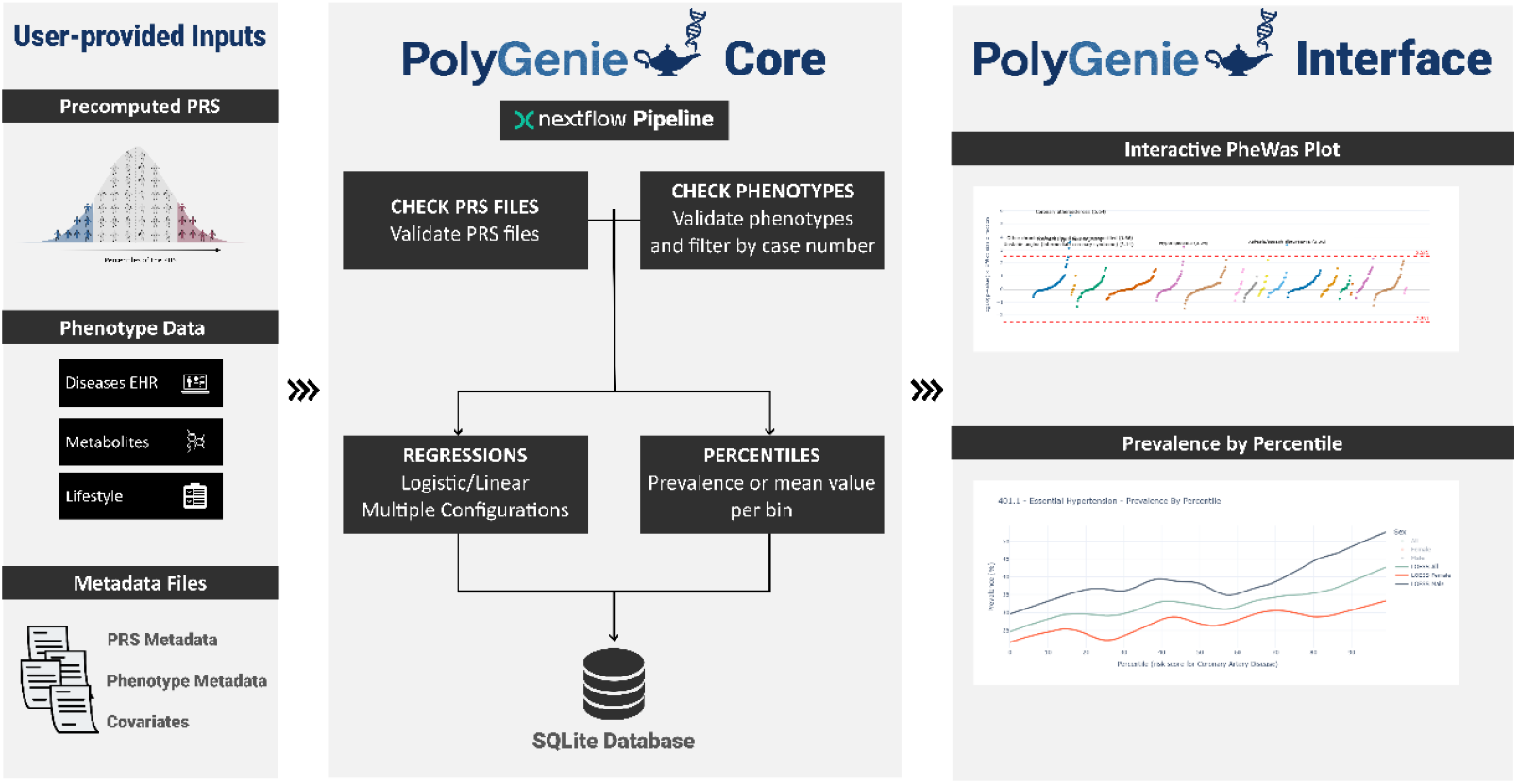
Overview of the PolyGenie pipeline. Left Panel: User-provided inputs including precomputed PRS and cohort phenotype data-Middle panel: Including preprocessing, PheWAS regression, and percentile analysis to a SQLite database. Right panel: representative visualization outputs from the web application, including the PheWAS scatter plot displaying signed log_10_(p-value) across phenotypic domains, and the percentile-based prevalence plot showing how outcome prevalence varies with genetic liability, stratified by sex.

**Figure 2.**
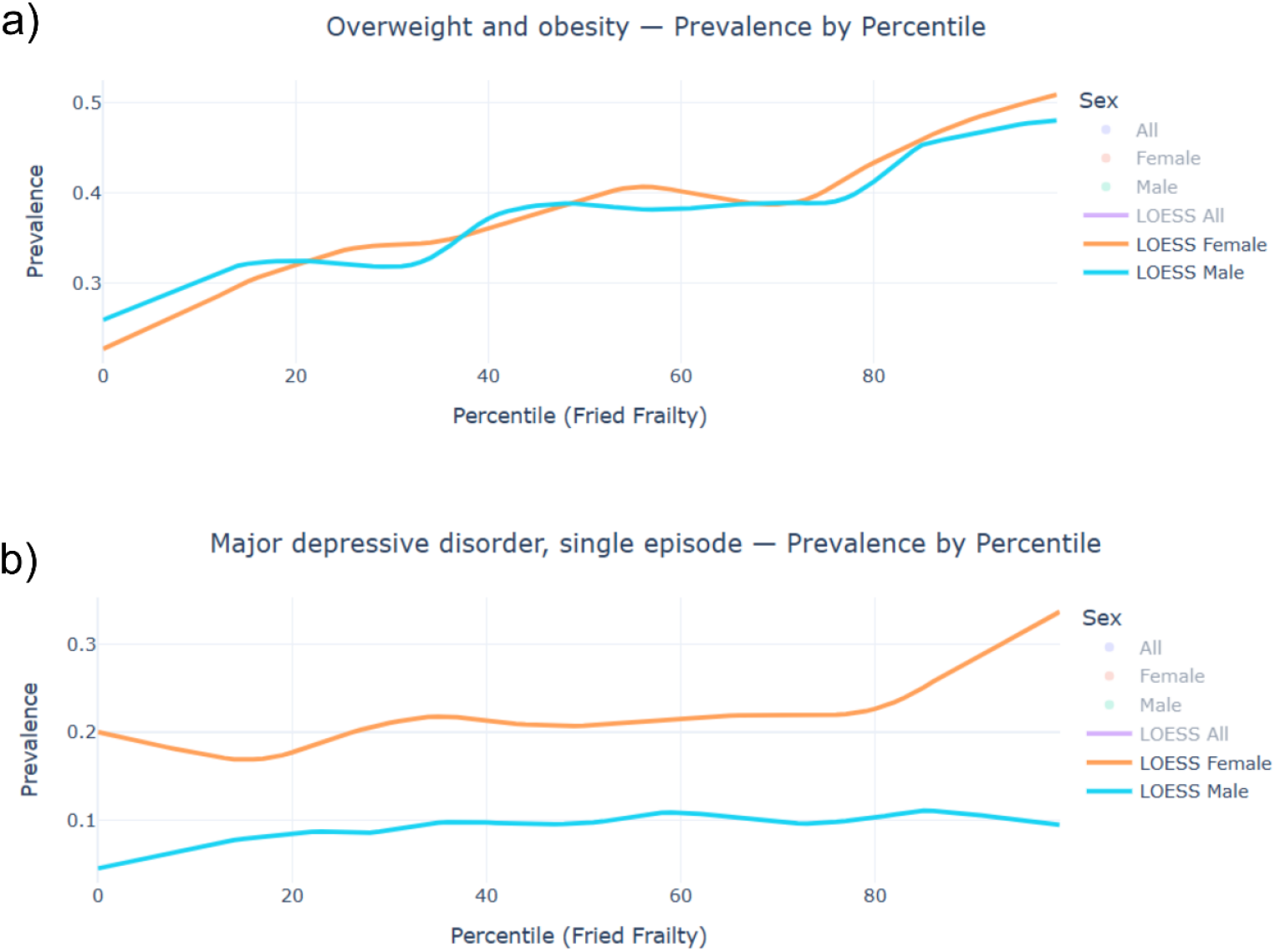
Prevalence of Clinical Outcomes Across Frailty PRS Percentiles. a) Prevalence of overweight and obesity (ICD-10: E66) increases approximately linearly with frailty polygenic risk score (PRS) percentile, with a comparable trend across sexes. b) Prevalence of major depressive disorder (ICD-10: F32) also rises with higher frailty PRS, with notably higher rates observed among females across all percentiles. These findings highlight the pleiotropic nature of genetic liability to frailty and its relationship with both metabolic and psychiatric conditions.

This example illustrates how PolyGenie enables rapid, hypothesis-generating exploration of pleiotropy and sex-specific genetic risk patterns within a harmonized cohort, without requiring custom analysis code.

## Conclusion

PolyGenie addresses a practical gap in the PheWAS tooling landscape by providing a portable, end-to-end pipeline that separates PRS computation from downstream analysis and visualization. Its Nextflow implementation ensures that the same pipeline runs on laptops, HPC clusters, and cloud environments, and its metadata-driven configuration requires no code changes when adapting to a new cohort. By accepting precomputed PRS from any method, it remains agnostic to the rapidly evolving landscape of PRS approaches, allowing users to apply best-practice scores without modifying the analysis framework.

Compared to existing resources, PolyGenie occupies a distinct niche within the PheWAS landscape. Static atlas-style platforms such as the PRS Atlas (5), SpPGS (6), and the PheWAS Catalog (11) provide valuable references for population-level associations but are inherently tied to their source data and not designed for redeployment on new cohorts. Analysis-focused tools such as the R PheWAS package (7) and pyPheWAS (8) offer computational flexibility but require programming expertise and lack integrated visualization pipelines. Specialized visualization platforms such as PheWAS-ME (9) and pyPheWAS Explorer (10) provide rich interactive dashboards but operate as post-hoc layers requiring prior external statistical analyses. PolyGenie bridges these fragmented approaches by providing a unified, end-to-end solution: it automates statistical analysis across hundreds of PRS-phenotype combinations, integrates results directly into an interactive database-backed web application, and remains portable and redeployable across arbitrary population cohorts through its metadata-driven design. This integration eliminates the operational burden of coordinating multiple specialized tools, enabling researchers to conduct systematic PheWAS analyses without programming expertise or manual result processing.

By enabling systematic cross-trait comparisons, generating testable hypotheses, and uncovering potential biomarkers or pleiotropic effects, PolyGenie serves both research and translational goals. It supports precision medicine by making complex genetic information accessible through user-friendly visual analytics. Developed under open data principles, aligned with ELIXIR interoperability standards, and hosted within a robust, publicly maintained infrastructure, PolyGenie is designed for long-term sustainability as a FAIR-compliant resource supporting durable genomic research across European cohorts.

The current implementation has limitations worth noting. The pipeline performs PheWAS in the frequentist regression framework and does not currently integrate Mendelian randomization or colocalization methods. The web application is optimized for exploration and hypothesis generation rather than for confirmatory statistical analysis. Extension to federated or multi-cohort analyses would require additional harmonization infrastructure beyond the current scope. These represent natural directions for future development.

Overall, PolyGenie provides a standardized, reproducible, and user-friendly framework for PheWAS analysis that can be readily deployed across diverse population cohorts, supporting systematic exploration of the shared genetic architecture of complex traits.

## Key Points

- PolyGenie is an open-source Nextflow pipeline for phenome-wide association studies (PheWAS) using polygenic risk scores, designed to be deployed on any cohort with minimal configuration.
- The pipeline accepts precomputed PRS from any method as input, separating PRS generation from downstream analysis and remaining agnostic to the choice of scoring approach.
- PolyGenie produces regression results and percentile-based prevalence estimates stored in a SQLite database, visualized through an interactive Dash web application with PheWAS plots and sex-stratified prevalence curves.
- A fully worked example using the GCAT cohort, applying over 100 PRS to clinical, molecular, and lifestyle phenotypes, is provided as a reproducibility and demonstration resource.
- The Nextflow implementation ensures portability across HPC, cloud, and local environments, supporting scalable parallel execution across large numbers of PRS and phenotypes.

## Availability and Implementation

PolyGenie is freely available at https://github.com/gcatbiobank/polygenie-pipeline under the MIT License. The repository includes full documentation, format specifications, and a demo dataset for local testing. The GCAT implementation is publicly accessible at https://polygenie.igtp.cat, and the modified version of the Dash interface at https://github.com/gcatbiobank/polygenie-gcat.

The pipeline is implemented in Nextflow DSL2 with Python analysis scripts. The web application uses the Plotly Dash framework with a SQLite backend. The GCAT deployment is hosted on infrastructure maintained by the Institut Germans Trias i Pujol and will remain publicly accessible for a minimum of two years following publication.

## Authors’ contributions

All authors meet the criteria for authorship. RdC and XF conceived and designed the study. MG developed the original analysis pipeline. XF adapted and extended the pipeline to Nextflow for community-wide applicability. NB led the implementation of the PheWAS analyses. RdC secured funding for the project. XF drafted the manuscript, with contributions from RdC and NB to the interpretation of results and overall framing of the work. All authors critically reviewed the manuscript for important intellectual content, approved the final version for publication, and agree to be accountable for all aspects of the work, ensuring that any questions related to its accuracy or integrity are appropriately investigated and resolved.

## Acknowledgments

We express our sincere gratitude to all GCAT cohort study participants for their invaluable contributions. We also thank the team at the Blood and Tissue Bank, coordinated by Dr. Alonso Nogués and Dr. Grífols, for their essential support in sample collection. Special thanks are extended to the GCAT project investigators for their ongoing commitment and collaboration. Electronic Health Record (EHR) data were provided by the Catalan Agency for Quality and Health Assessment (PADRIS Program). A complete list of GCAT investigators can be found at www.genomesforlife.com.

